# Reconstructed human pigmented skin/epidermis models achieve epidermal pigmentation through melanocore transfer

**DOI:** 10.1101/2022.03.03.482849

**Authors:** Michael J. Hall, Sara Lopes-Ventura, Matilde V. Neto, João Charneca, Patricia Zoio, Miguel C. Seabra, Abel Oliva, Duarte C. Barral

## Abstract

The skin acts as a barrier to environmental insults and provides many vital functions. One of these is to shield DNA from harmful UV radiation, which is achieved by skin pigmentation arising as melanin is produced and dispersed within the epidermal layer. This is a crucial defence against DNA damage, photo-ageing and skin cancer. The mechanisms and regulation of melanogenesis and melanin transfer involve extensive crosstalk between melanocytes and keratinocytes in the epidermis, as well as fibroblasts in the dermal layer. Although the predominant mechanism of melanin transfer continues to be debated and several plausible models have been proposed, we and others previously provided evidence for a coupled exo/phagocytosis of melanocore model. Herein, we performed histology and immunohistochemistry analyses and demonstrated that a newly developed full-thickness 3D reconstructed human pigmented skin model and an epidermis-only model exhibit dispersed pigment throughout keratinocytes in the epidermis. Transmission electron microscopy revealed melanocores between melanocytes and keratinocytes, suggesting that melanin is transferred through coupled exocytosis/phagocytosis of the melanosome core similar to our previous observations in human skin biopsies. We therefore present evidence that our *in vitro* models of pigmented human skin show epidermal pigmentation comparable to human skin. These findings have a high value for studies of skin pigmentation mechanisms and pigmentary disorders, whilst reducing the reliance on animal models and human skin biopsies.

**Significance:** Currently, many approaches to study skin pigmentation and pigmentary disorders use 2D cultures or co-culture of melanocytes and/or keratinocytes, as well as skin biopsies obtained from consenting donors. However, challenges are faced when it is necessary to mimic the tissue microenvironment as there is a limited availability of skin biopsies, especially for the case of rare pigmentary disorders. Through this study, we show the potential of 3D *in vitro* reconstructed pigmented skin/epidermis models for both general and specific pigmentary research questions, which could reduce reliance on animal models and human skin biopsies.

## Introduction

The skin is the largest organ of the human body and is the first-line defence against environmental insult and pathogens. Vital roles of the skin include protection against UV radiation (UVr), thermoregulation and infection prevention (Roger et al., 2019; Chambers & Vukmanovic‐Stejic, 2020). The superficial-most skin structure comprises 2 main layers - the dermis and the epidermis. The dermis measures 3-4 mm in thickness and is mostly composed of fibroblasts, which produce the extracellular matrix to support the epidermis and impart strength to the tissue. The epidermis is the outermost layer of the skin, predominantly composed of keratinocytes in four different stages of differentiation that are referred basally to apically as *stratum basale, stratum spinosum, stratum granulosum and stratum corneum*, respectively (Roger et al., 2019; Chambers & Vukmanovic‐Stejic, 2020). The *stratum basale* is responsible for maintaining the constant renewal of epidermal cells, as here resides a layer of undifferentiated keratinocytes. In the *stratum spinosum*, cells begin to differ in morphology and produce different keratins from the basal layer (Nestle et al., 2009). Above this layer, the keratinocytes acquire a more flattened shape until reaching the *stratum granulosum*, where keratin synthesis increases (Mehrel, 1990). The *stratum corneum* is the outermost layer and is mostly responsible for barrier function of the skin (Nestle et al., 2009).

The epidermis also comprises melanocytes in the basal layer, which extend dendrites to contact up to 40 keratinocytes (Hoath & Leahy, 2003). To achieve epidermal pigmentation, melanin is synthesised and packaged by melanocytes in the form of lysosome-related organelles termed melanosomes, before the melanin is secreted from melanocyte dendrite tips (Chambers & Vukmanovic‐Stejic, 2020). The melanin is subsequently transferred to adjacent keratinocytes, where it is stored in compartments that together form a supra-nuclear ‘cap’ to shield DNA from UVr (Benito-Martínez, 2021). The resulting skin pigmentation exhibits a photoprotective effect against UV-induced damage of cells, protection against photo-ageing, and skin cancer (Lin & Fisher, 2007; Zhang & Duan, 2018). Highlighting the importance of photoprotection, epidemiological studies demonstrate that fair skin individuals are 20-70 times more likely to develop skin cancer than dark skin individuals (d’Ischia et al., 2015).

Melanocyte-keratinocyte communication is vital for control of skin pigmentation, which lends to the term ‘epidermal-melanin unit’ to refer to the intercellular crosstalk modulating melanogenesis and melanin transfer. For example, photoprotection can be enhanced by UVr exposure in a process known as adaptive skin pigmentation, or more commonly as “tanning”. In this process, keratinocytes secrete soluble ɑ-melanocyte stimulating hormone (ɑ-MSH) upon exposure to UVr, which stimulates melanogenesis by activating melanocortin-1 receptor (MC1R) on melanocytes and increasing microphtalmia-associated transcription factor (MITF) levels (Lin & Fisher, 2007). Furthermore, other factors such as endothelin-1 (EDN1) and membrane-type stem cell factor (SCF) contribute to autocrine and paracrine feedback loops to sustain the increase in MITF (Imokawa, 2004), and exosomes secreted by keratinocytes have been shown to increase melanogenesis by melanocytes (Lo Cicero et al., 2015). Additionally, factors produced by fibroblasts in the dermis such as soluble type SCF and hepatocyte growth factor (HGF) can also stimulate melanocytes. Dysfunction of these feedback loops and signalling pathways is linked with several pigmentary disorders (Imokawa, 2004).

The mechanism(s) of melanin transfer from melanocytes to keratinocytes continue to be debated, with several models proposed and evidenced in the literature. Four main models currently exist, reviewed recently by us (Moreiras et al., 2021a) and others (Benito-Martínez et al., 2021). Briefly, these are: coupled exo/endocytosis of the melanosome core; cytophagocytosis of melanocyte dendrites; fusion of melanocyte and keratinocyte membranes to allow direct transfer of melanosomes; and shedding of melanosome-laden vesicles/globules by melanocytes. Our lab previously published evidence showing naked melanin between cells in human skin biopsies (Tarafder et al., 2014), and we demonstrated that Rab11b and the exocyst complex are required for melanocore exocytosis from a melanocyte cell line by fusion of melanosomes with the plasma membrane (Tarafder et al., 2014; Moreiras et al., 2019), both of which support the exo/endocytosis model. Furthermore, we have demonstrated that the melanosome core, or melanocore, uptake by keratinocytes occurs through phagocytosis rather than macropinocytosis (Moreiras at al., 2021b), before melanin is stored within a non-degradative compartment in keratinocytes with a single limiting membrane (Correia et al., 2018; Hurbain et al., 2018). We recently proposed the term ‘melanokerasome’ (MKS) to refer to this melanin-containing compartment within keratinocytes, to distinguish it from melanosomes in melanocytes. Although different transfer mechanisms could co-exist in physiological and pathological circumstances, mounting evidence supports the model of exo/phagocytosis of melanocores as a major mechanism in constitutive skin pigmentation.

Reliable and replicable models to study the control of skin pigmentation are of high value for drug development, disease modelling and basic discoveries, but existing models have drawbacks. For example, monoculture systems lack the three-dimensional structure and variable keratinocyte differentiation state of stratified skin epidermis, and also do not recapitulate the aforementioned intricate crosstalk that exists between the cells of the epidermis and dermis. Skin biopsies and explants are also utilised in some studies, although their availability and long-term viability in culture is restrictive. Additionally, although many discoveries of mechanisms of skin pigmentation were made using rodents, it should be noted that the most utilised animal model, the mouse, lacks pigment in skin and there is mounting opposition to the use of animals in the cosmetic and scientific industries. In recent publications, we described a model of full-thickness human skin formed using primary human keratinocytes and fibroblasts cultured on a fibroblast-derived matrix (Zoio et al., 2021a) that can also be formed on a chip (Zoio et al., 2021b) for real-time barrier function analysis. These artificial skins can be maintained with an air-liquid interface (ALI) for up to 50 days and exhibit the stratified layers of epidermis as seen in human skin *in vivo*.

Herein, we characterise the epidermal pigmentation of the newly developed reconstructed human pigmented skin (RHPS) model, and present evidence of melanocore transfer from melanocytes to keratinocytes. In addition, we present a comparable reconstructed human pigmented epidermis (RHPE) model that can be formed in a more rapid and simple manner, and with fewer resources required.

## Materials and Methods

### Cell Culture

For the RHPS/RHPE cultures the following primary cells were used: neonatal human dermal fibroblasts (HDFn, Innoprot P10857, used for RHPS only), neonatal human epidermal melanocytes from darkly pigmented skin (HEM-DP, GIBCO), and neonatal human epidermal keratinocytes (HEK, GIBCO). HDFn were cultured in high glucose Dulbecco’s Modified Eagle medium (DMEM, Gibco) supplemented with 10% fetal bovine serum (FBS, Gibco) and 1% penicillin-streptomycin solution (pen-strep, Gibco). HEM-DP were cultured in melanocyte medium 254 (M254, Gibco) supplemented with 1% human melanocyte growth supplement (HMGS, Gibco) and 1% Pen-strep. The cells were passaged at a 1:2 ratio when they reached 90% of confluency. HEK were cultured in EpiLife medium (Gibco), supplemented with 0.6 mM CaCl_2_ (Gibco), 1% human keratinocyte growth supplement (HKGS, Gibco) and 1% pen-strep. HDFn and HEM-DP were used until passage 13 and HEK until passage 7 (for RHPS) or passage 4 (for RHPE). All cells were maintained at 37°C and 5% carbon dioxide in a humidified incubator.

For RHPS, a polystyrene scaffold (Alvetex®, REPROCELL Europe Ltd., UK, #AVP005-48) with an area of 1.13 cm^2^ was used to develop the dermis, and a well insert with a 0.78 cm^2^ polycarbonate membrane (Merck Millipore, #PIHP01250) coated with bovine collagen I (Gibco) was used in RHPE, both in 6-well plates (one scaffold or membrane per well). The models were formed and maintained at 37°C and 5% carbon dioxide in a humidified incubator. RHPS and RHPE were developed according to the protocol already published (Zoio et al. 2021a) and the formation of both models is detailed and compared in results, and summarised in Figure 1. The cells used were from commercial sources, and thus studies are exempt from IRB review.

**Figure 1:**
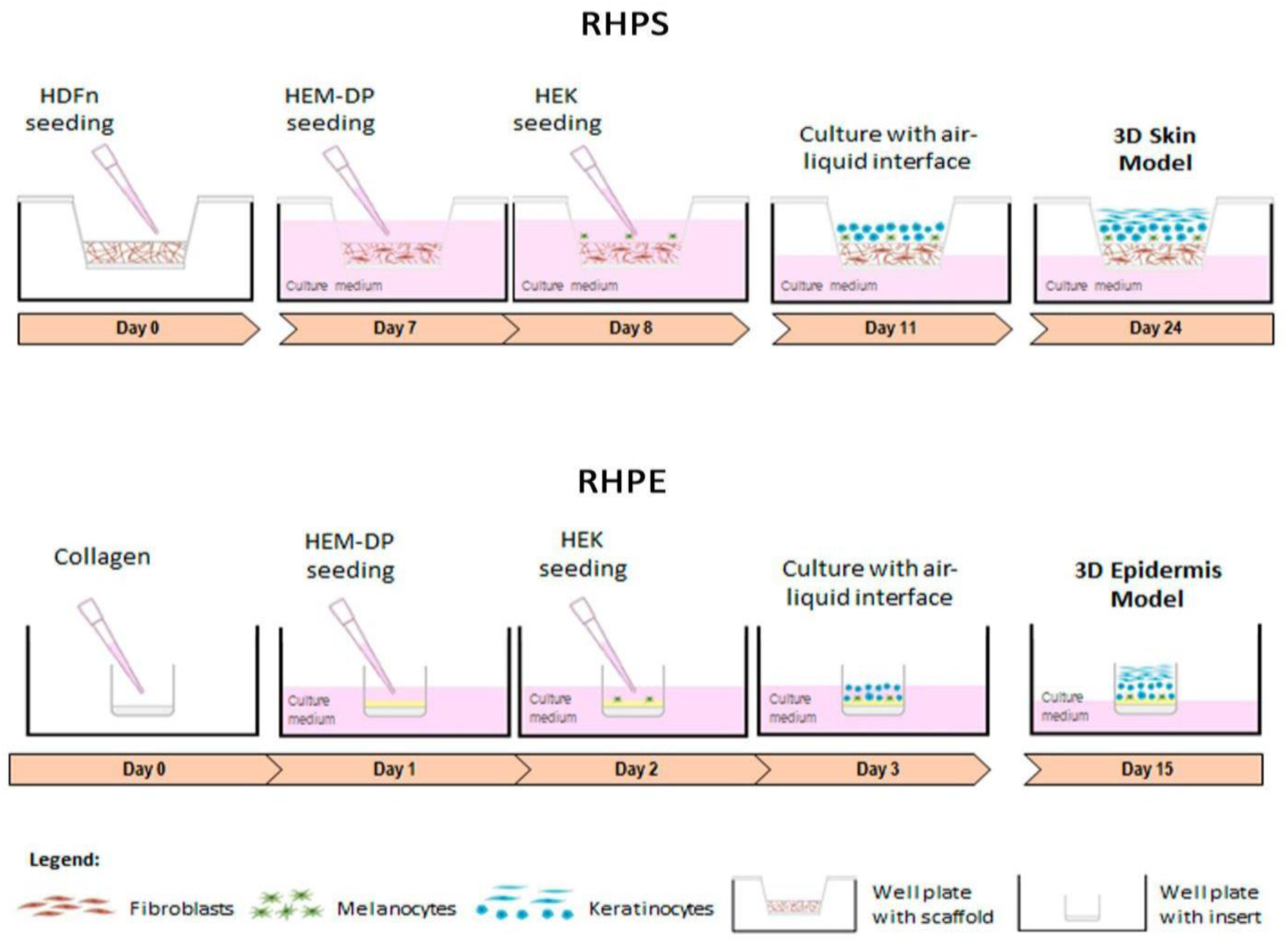
Schematic representation depicting the stages of forming reconstructed human pigmented skin and epidermis. RHPS: On day 0, fibroblasts (HDFn) are seeded onto the scaffold to allow the formation of the dermis. On day 7, the formation of the epidermis begins with the addition of melanocytes (HEM-DP) followed by addition of keratinocytes (HEK) on the following day. Three days after, apical media is removed to achieve the air-liquid interface and on day 24, the model is fully formed. RHPE: The epidermis model is formed on a collagen-coated membrane to support melanocytes and keratinocytes. On day 1, the melanocytes are seeded, followed by keratinocytes on the following day. One day after, the air-liquid interface is established and on day 15 the epidermis is completed.

### Histology and Immunohistochemistry

RHPS or RHPE models were fixed in 10% neutral buffered formalin (Bio-Optica). The samples were dehydrated, paraffin-embedded and sections of 5 μm thickness were prepared for staining with hematoxylin and eosin (H&E) and Fontana-Masson.

For immunohistochemical analysis, 5 µm thick sections of the RHPS/RHPEs were mounted on glass slides. These were stained with anti S-100 (polyclonal, Dako) or anti-tyrosinase (T311, Novocastra) primary antibodies diluted in phosphate buffered saline containing 1% bovine serum albumin (PBS-BSA 1%; BSA, Sigma).

### Transmission Electron Microscopy (TEM)

All reagents and materials were purchased from Electron Microscopy Sciences unless otherwise stated. Specimens were fixed in 2% paraformaldehyde, 2% glutaraldehyde in 0.1 M phosphate buffer (PB) at pH 7.4 overnight at 4⁰C. After washing with PB, specimens were post-fixed for 1 hour on ice using 1% osmium tetroxide and 1.5% potassium ferrocyanide in distilled water. Specimens were subsequently dehydrated with a series of increasing ethanol concentrations (35%, 45%, 50%, 70%, 90%, 2x 100%) before infiltrating and embedding in Epon resin. After polymerising at 70⁰C overnight, resin blocks were sectioned at 90-100 nm using a UC7 ultramicrotome (Leica) and a diamond knife (Diatome), and sections collected on formvar/carbon-coated copper mesh grids. Sections on grids were post-stained with lead citrate and imaged using a Hitachi H-7650 TEM equipped with an AMT XR41M digital camera, or an FEI Tecnai G2 Spirit BioTWIN TEM equipped with an Olympus-SIS Veleta CCD camera.

One specimen of RHPS and three specimens of RHPE were processed and observed by TEM. Quantitative TEM analysis was performed on a single specimen of RHPE, which was representative of the three biological replicates. Melanocytes and keratinocytes were identified and distinguished using criteria listed in Table 1.

**Table 1:** Criteria used for distinguishing melanocytes and keratinocytes by transmission electron microscopy. Features used to correctly identify melanocytes and keratinocytes by TEM are listed.

| Feature | Melanocytes | Keratinocytes |
| --- | --- | --- |
| **Cytoskeletal features** | Not clearly observed | Bundles of actin/keratin fibrils often visible |
| **Desmosomes** | Absent between melanocytes and keratinocytes | Present between adjacent keratinocytes, predominantly in *stratum spinosum* |
| **Electron density** | Relatively electron-dense cytoplasm | Relatively electron-lucent cytoplasm, especially in apical layers |
| **Glycogen granules** | Few cytoplasmic glycogen granules | Many cytoplasmic glycogen granules |
| **Melanin content** | Densely packed with melanosomes | Sparsely packed with melanokerasomes (MKS) |
| **Position** | Cell body in the basal layer only | In all the stratified layers |
| **Shape** | Usually visible as small cylindrical dendrites, except when cell body is observed in *stratum basale* | Large, flat cells often with flattened nucleus |

## Results

### Reconstructed human pigmented skin and epidermis models are histologically comparable to human skin

A schematic representation of the production timeline of RHPS and RHPE models is presented in Figure 1, based on a protocol previously developed by us (Zoio et al., 2021a). The RHPS model constitutes three primary human cell types from commercial sources: melanocytes, keratinocytes and fibroblasts, whereas the RHPE model omits the latter. The RHPS models are formed with a total of 1×10^6^ fibroblasts, 5×10^5^ keratinocytes and 5×10^4^ melanocytes. A 1:10 ratio of melanocytes to keratinocytes is used in both RHPS and RHPE. To form the RHPS model, a scaffold is used to support fibroblasts, which spontaneously produce a matrix to recreate the dermis (Figure 1a, Figure S1). The number of fibroblasts was previously optimised to originate a fully mature structure capable of supporting the epidermis and avoiding keratinocyte infiltration into the dermal compartment (Zoio *et al*., 2021a). On the other hand, in the absence of the dermis layer, a collagen-coated polycarbonate membrane is used to support the RHPE model (Figure 1b). For this model, 3×10^4^ cells/mL of melanocytes are seeded upon the collagen layer, which was laid 24 h earlier, and after 1 day, 3×10^5^ cells/mL of keratinocytes are seeded. Both models form an epidermis layer, which involves the introduction of melanocytes in the basal lamina, followed by the addition of keratinocytes the following day (Figure 1). The removal of the apical media creates an ALI that induces keratinocyte stratification and confirms the integrity and barrier functions of the RHPS and RHPE models. Although the RHPE model lacks the recreated dermis layer, its formation can be completed in around 9 days fewer than the RHPS model.

We previously demonstrated that the RHPS model shows expression of typical epidermal markers, indicating proper proliferation and differentiation of keratinocytes (Zoio et al. 2021a). In this model, fibroblasts produce their own extracellular matrix (ECM) in the absence of materials of animal origin to recreate the dermis (Figure 2a, Figure S1) and the keratinocytes produce a stratified and differentiated epidermis in both models (Figure 2a and c, Figure S2). The ECM and the fibroblasts incorporated within the scaffold form the model dermis.

**Figure 2:**
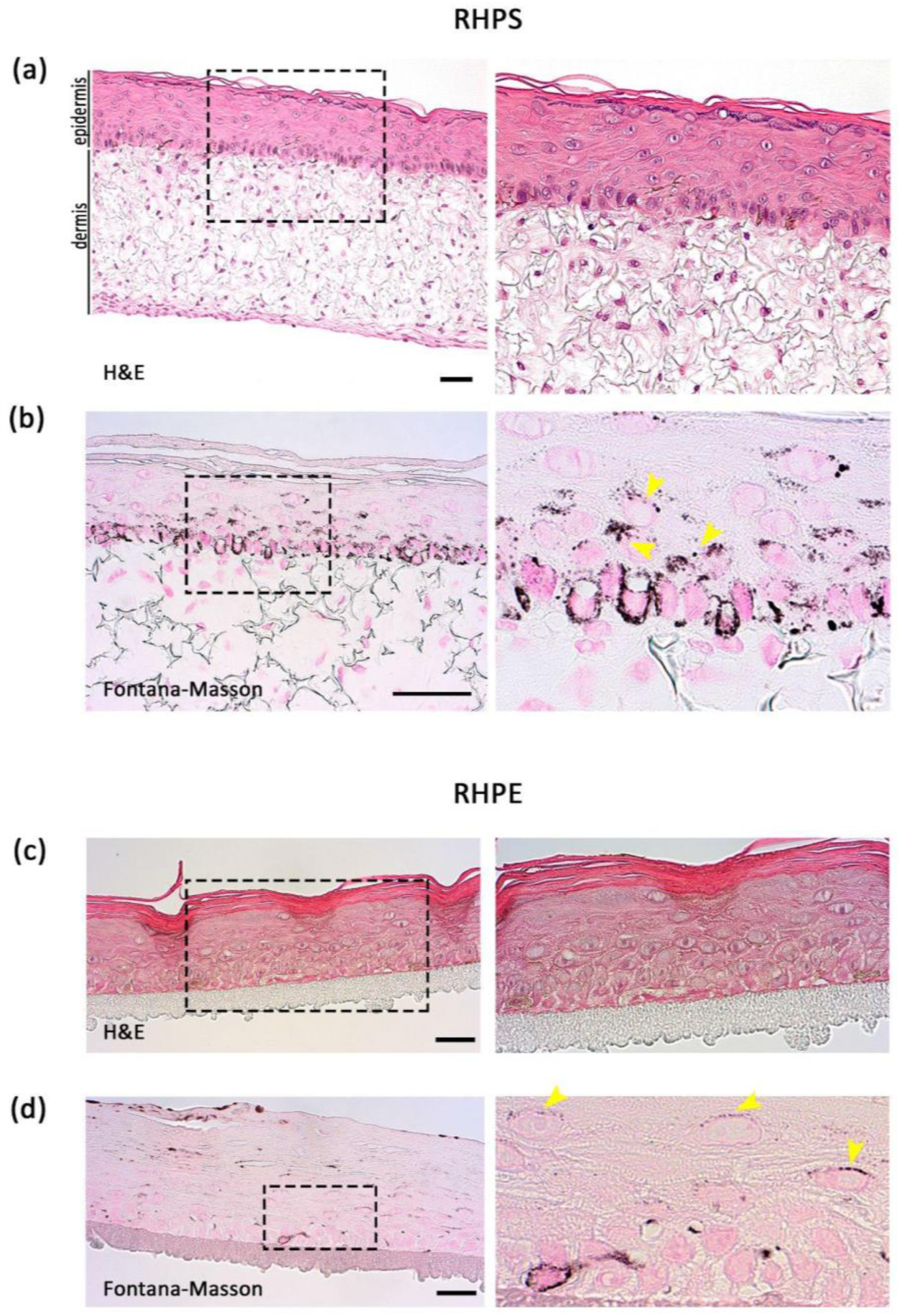
Histology of reconstructed human pigmented skin and epidermis models. RHPS **(a)** and **(b)**, RHPE **(c)** and **(d)**. Haematoxylin-Eosin (H&E) staining is shown in **(a)** and **(c)**, and Fontana-Masson staining is shown in **(b)** and **(d)** to identify pigment in melanocytes and keratinocytes. Dermis is visible in the skin model and the stratification and differentiation of epidermis are visible in both models. Magnified views of the boxed areas are shown in the right panels. Images are representative of three independent experiments. Arrowheads indicate areas of pigment accumulation in keratinocytes. Scale bars, 50 μm.

### Reconstructed human pigmented skin and epidermis models show pigment dispersion throughout the epidermis

We next aimed to characterise melanocyte distribution and melanin dispersion in our models. Pigment in the epidermis was identified with Fontana-Masson staining (Figure 2b and d), and melanocytes were identified by immunohistochemistry of two melanocyte-specific proteins - tyrosinase and S100. Melanocyte alignment comparable to human epidermis *in vivo* was observed in the basement membrane in RHPS (Figure 3a and b) and in RHPE models (Figure 3c and d). In both models, melanin transfer is functional throughout the epidermis, evidenced by melanin within keratinocytes, highlighted with yellow arrows in the right panel of b and the right panel of d in Figure 2.

**Figure 3:**
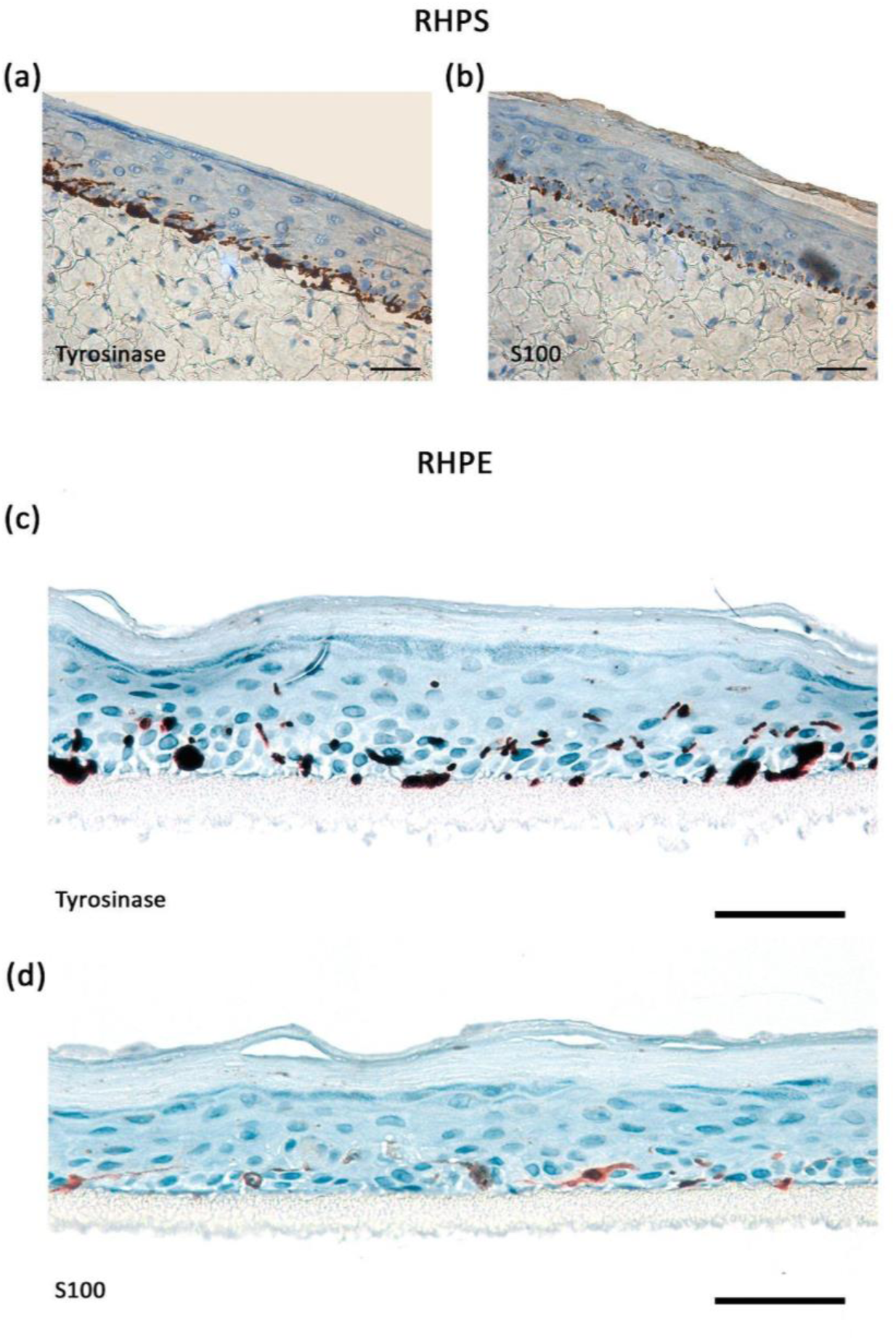
Immunohistochemical analysis of epidermal melanocytes in reconstructed human pigmented skin and reconstructed human pigmented epidermis models. Melanocytes in the basal layer were identified by staining for tyrosinase or S100 proteins in RHPS **(a)** and **(b)** and RHPE **(c)** and **(d)**. Images are representative of three independent experiments. Scale bars, 50 μm.

It is estimated that in human skin, keratinocytes outnumber melanocytes in a ratio of around 1 melanocyte to every 40 keratinocytes (Hoath & Leahy, 2003). Thus, a small number of melanocytes must supply melanin to a relatively large number of keratinocytes. A resulting feature of melanocytes is the development of actin-rich dendrites that extend from the cell body to reach multiple keratinocytes (Fitzpatrick, 1967). Melanin is transported along and secreted from these dendrites, which facilitates efficient pigment dispersion throughout multiple keratinocytes in the epidermis (Tian et al., 2020). We performed transmission electron microscopy (TEM) on specimens of RHPS and RHPE to reveal features that facilitate the efficient pigment dispersal in our models. Keratinocytes were distinguished from melanocytes by identifying contrasting hallmarks of each cell type, which are summarised in Table 1. For example, keratinocytes often present with desmosomes at their interface with adjacent keratinocytes, contain more glycogen granules, and often appear larger and flatter than the melanocyte dendrites. On the other hand, whilst only few melanocyte cell bodies are observed in ultrathin sections of RHPS and RHPE by TEM (such as in Figure 4a), melanocyte dendrites are very often identified between keratinocytes by the presence of densely packed melanosomes (Figure 4b and 4c). This indicates that the extensive epidermal pigmentation observed in our models is at least in part due to the formation of extensive melanocyte dendrites as seen *in vivo*.

**Figure 4:**
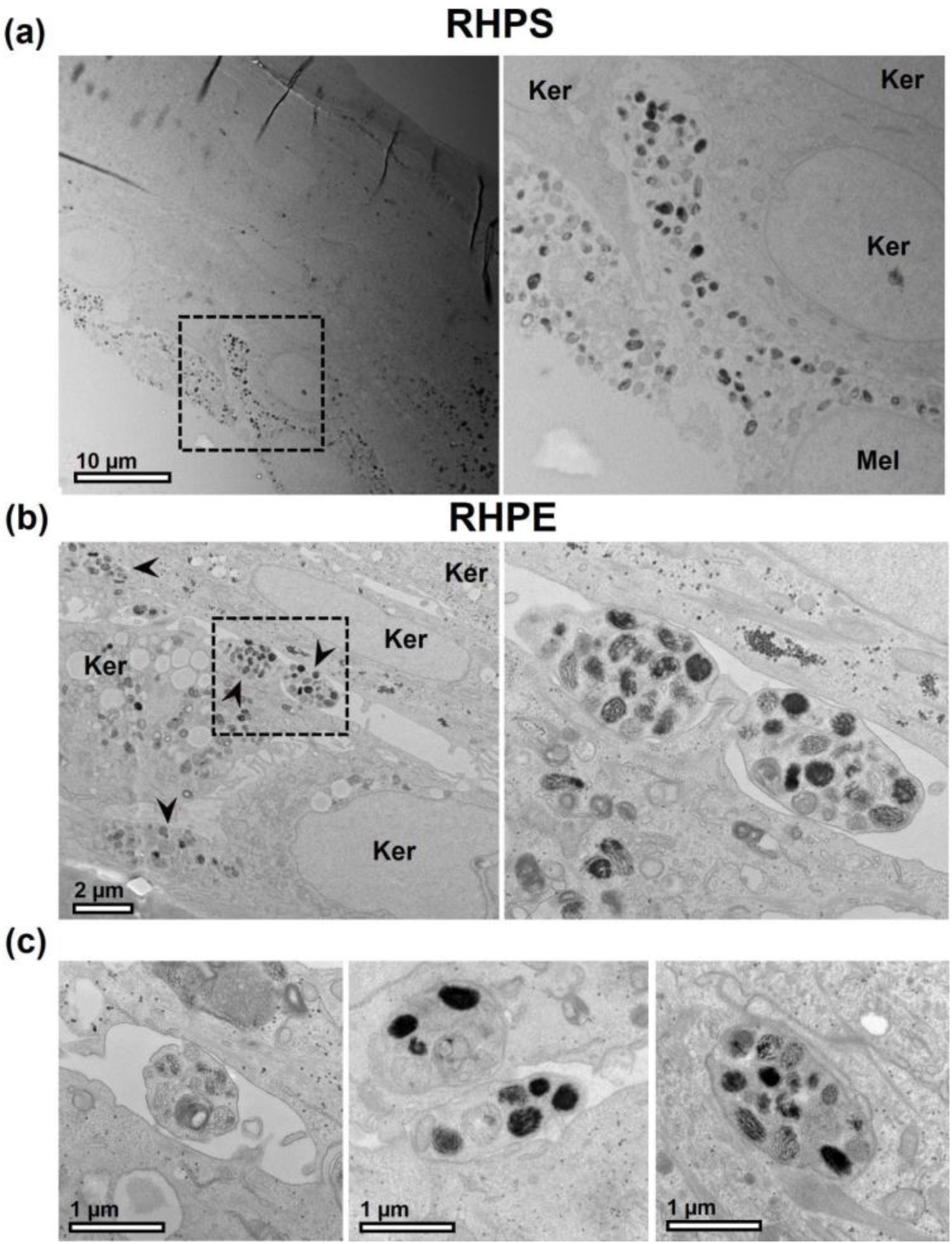
Melanocyte dendrites extending between keratinocytes. RHPS **(a)** or RHPE **(b)** and **(c)** were observed by transmission electron microscopy. Multiple dendrites extending from the cell bodies of melanocytes (Mel) are often visible between keratinocytes (Ker), and are indicated by arrowheads in (b). Scale bar, 10 μm (a), 2 μm (b), 1 μm (c).

### Extracellular melanocores demonstrate exocytosis of melanosome content

Multiple models exist for the mechanism(s) of melanin transfer from melanocytes to keratinocytes (Moreiras et al. 2021a; Benito-Martínez et al., 2021). We previously demonstrated the presence of melanocores between the plasma membranes of melanocytes and keratinocytes in human skin biopsies (Tarafder et al., 2014). This is consistent with the exo/endocytosis model of melanin transfer, whereby the fusion of melanosomes with melanocyte plasma membrane releases melanin pigment that is taken up by adjacent keratinocytes, decorating the melanocores within a single membrane. Recently, we provided evidence that this internalisation of melanocores by keratinocytes occurs by phagocytosis (Moreiras et al. 2021b). Having observed by histology that our RHPS/RHPEs exhibit melanin in keratinocytes, which indicated functional pigmentation comparable to human skin *in vivo*, we next aimed to confirm that melanocores are transferred from melanocytes to keratinocytes in our models by identifying key features by TEM.

When focussing on the melanocyte dendrites in the models at higher magnification, examples of melanocores were observed in both RHPS (Figure 5a) and RHPE (Figures 5b and 5c). In Figure 5c i., a melanosome likely in the process of fusing with the plasma membrane is highlighted by the arrowhead, and a melanocore is observed between the melanocyte and keratinocyte plasma membranes. Figure 5c ii. shows a melanocore that has likely recently been transferred to the keratinocyte from the pictured melanocyte. In an approximately 500 µm length of an ultrathin section of RHPE, 20 melanocyte dendrites were clearly identified and quantitative analysis revealed that each dendrite contacted at least two keratinocytes. Six of these dendrites (representing 30%) presented with melanocores within a 0.5 µm distance of their plasma membranes in the section plane, suggesting that they originated from the melanocyte dendrite in the image. These observations suggest that melanocores are indeed secreted from melanocyte dendrites in these 3D models to efficiently disperse pigment, consistent with our previous observations in human skin (Tarafder et al., 2014).

**Figure 5:**
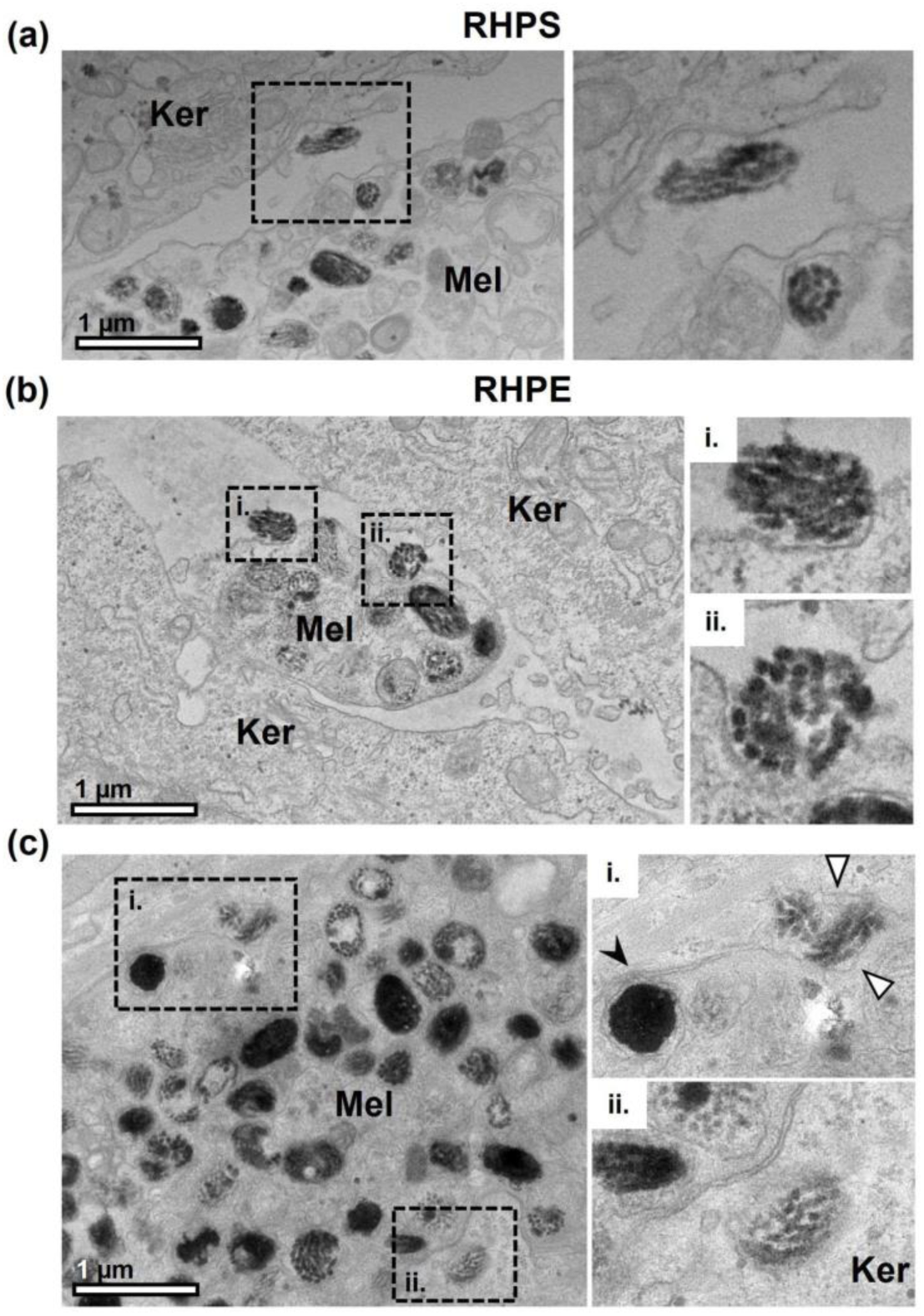
Melanocores in the extracellular space between melanocyte dendrites and keratinocytes. Melanocyte dendrites (Mel) are observed between keratinocytes (Ker) in RHPS **(a)** and RHPE **(b)** and **(c)** by TEM. The boxed regions are shown at higher magnification in the right panel. Several examples of melanocores between the cells and close to melanocyte dendrites are shown in (a), (b) and (c). A melanosome in the process of fusing with the melanocyte plasma membrane is visible in (c) i., indicated by the black arrowhead, and a melanocore is observed between the melanocyte and keratinocyte plasma membranes, indicated by the white arrowheads. In (c)ii., a recently transferred melanocore is observed in the keratinocyte. Scale bar, 1 μm.

### Melanocores are internalised by keratinocytes and are encased by a single membrane

According to the coupled exo/phagocytosis model of melanin transfer, melanocores are phagocytosed by adjacent keratinocytes, thus encasing them with a single membrane originating from the keratinocyte plasma membrane. However, to our knowledge, evidence of melanocore phagocytosis by TEM does not exist in the literature. In RHPE, we observed examples of melanocores that were in the process of being engulfed by keratinocytes, likely by phagocytosis (Figures 6a, 6b). The newly-formed single limiting membrane derived from the keratinocyte plasma membrane is highlighted by arrowheads in Figure 6a ii and 6b (inset). Thus, we believe this to be the first visualisation of melanocore internalisation by keratinocytes using TEM, which further supports the exo/phagocytosis model proposed by us.

**Figure 6:**
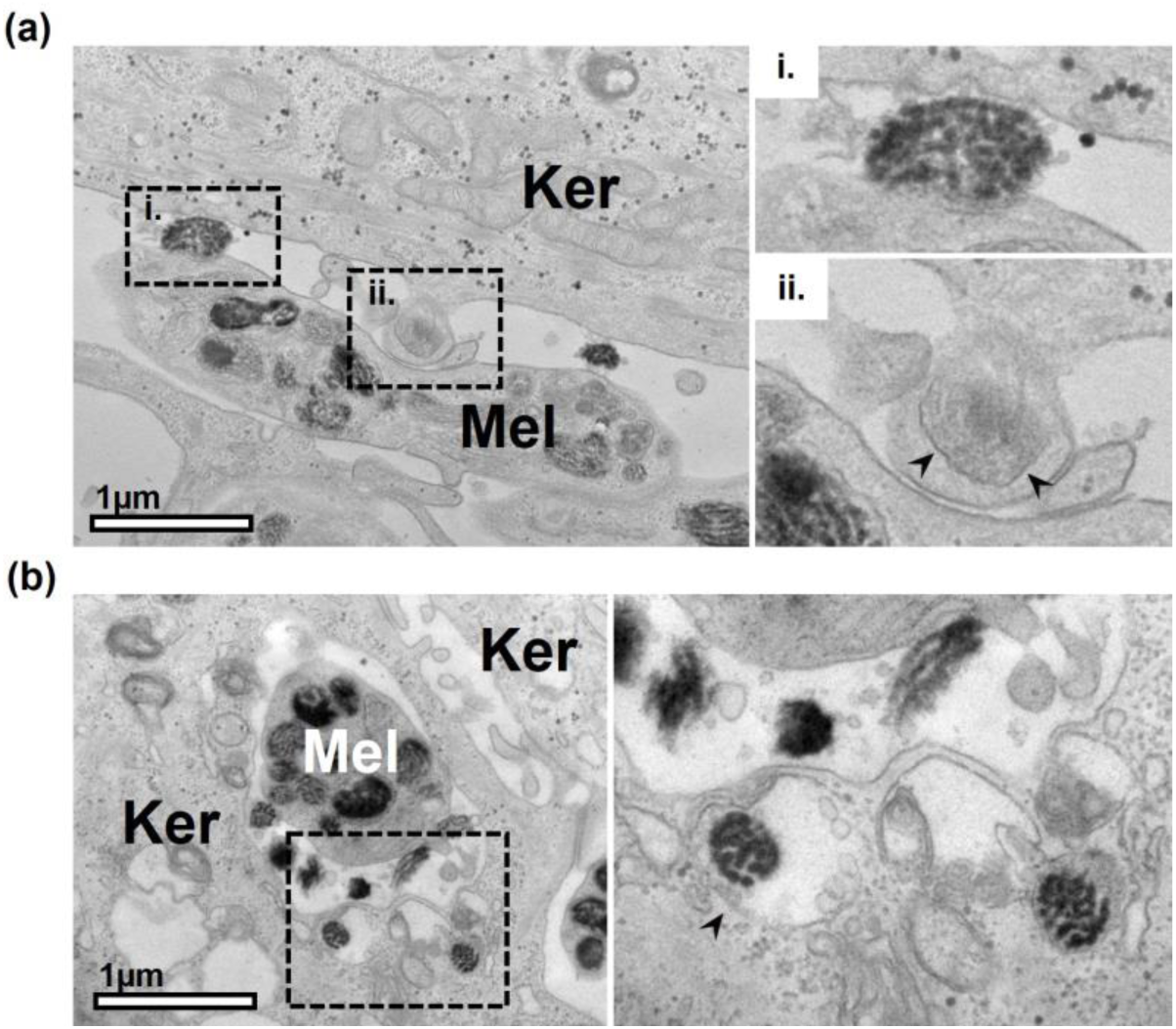
Melanocores in the process of engulfment by keratinocytes in reconstructed human pigmented epidermis. Examples of melanocyte dendrites (Mel) are observed between keratinocytes (Ker) in RHPE. In addition to melanocores in the extracellular space in (a) i. and (b), melanocores can be observed that have recently been engulfed by the adjacent keratinocytes [(a) ii and right panel of (b)]. Arrowheads indicate the membrane encasing the melanocore that is derived from the plasma membranes of the keratinocytes. Scale bar, 1 μm.

We next identified keratinocytes by TEM, again using criteria outlined in Table 1, to establish whether the melanin-containing compartments (melanokerasomes - MKSs) possess a single limiting membrane, as per the model of coupled exo/phagocytosis of melanocores. We observed that melanin is indeed surrounded by a single limiting membrane forming MKSs (Figure 7, black arrowheads). This is clearly contrasted with double membranes of mitochondria and the nuclear envelope (Figure 7, white arrowheads). Moreover, in a specimen of RHPE, we identified 15 keratinocytes and counted the number of limiting membranes present around all of their MKSs. Of the 71 MKSs observed within these keratinocytes, 64 were clearly encased by only a single limiting membrane around the majority of their perimeter (90%). The outlying 7 MKSs counted did not possess a clear single limiting membrane, and instead appeared to contain membrane whorls typical of lysosomes or resided within autophagosomes. We therefore propose that these are likely MKSs that have been sequestered by autophagy, rather than representing an alternative mode of internalisation. Thus, the appearance of melanin within keratinocytes in our models is consistent with the hypothesis that keratinocytes phagocytose melanocores as a major pigment transfer mechanism in the RHPS and RHPE models.

**Figure 7:**
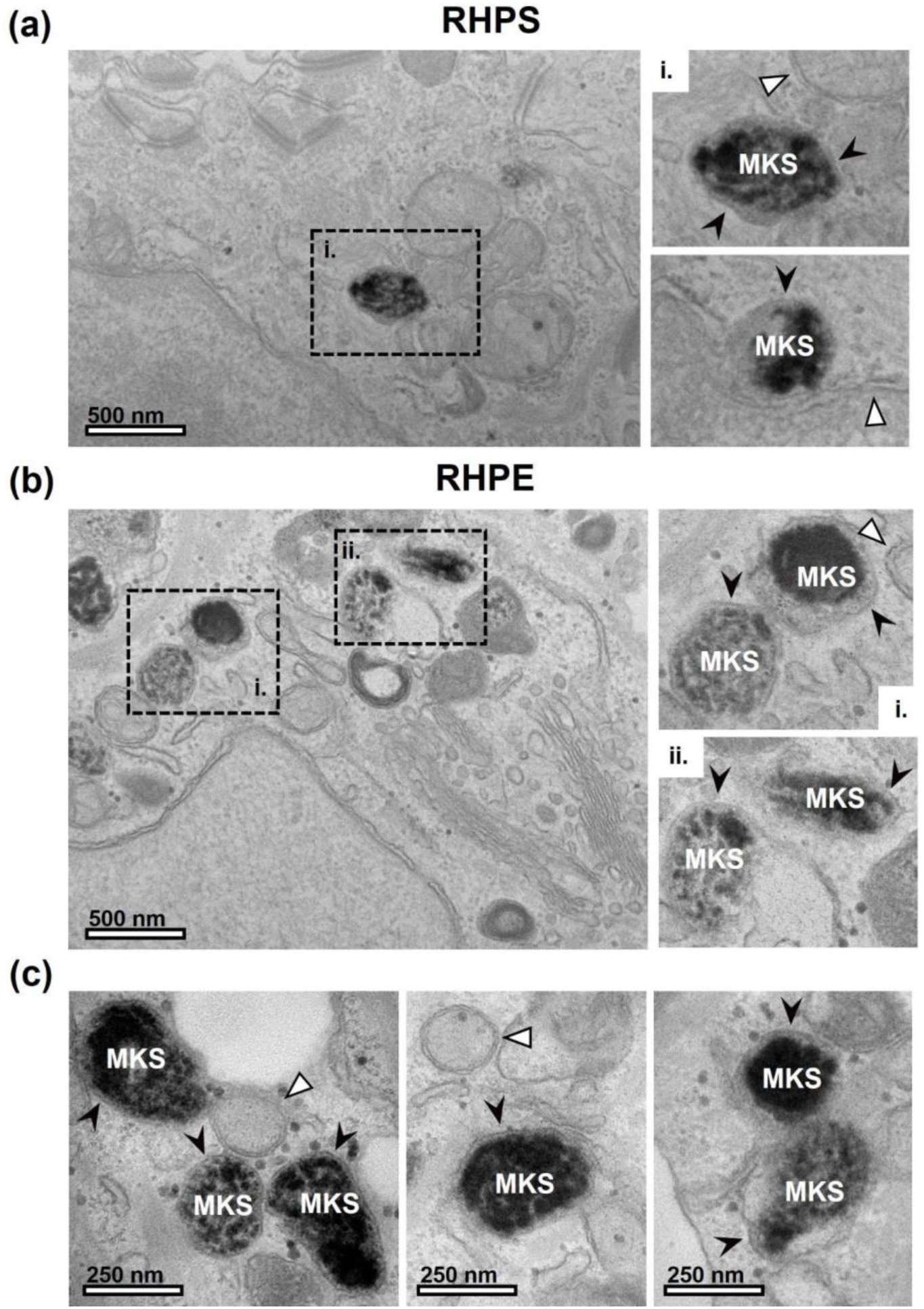
Melanin within keratinocytes is surrounded by a single membrane. In keratinocytes, melanin pigment is observed in a compartment surrounded by a single limiting membrane in RHPS **(a)** and RHPE **(b)** and **(c)**, which we refer to as melanokerasome (MKS). The boxed regions are shown in the right panel. In contrast, double membranes are clearly observed surrounding mitochondria and the nucleus (indicated by white arrowheads). Scale bar, 500 nm (a and b) or 250 nm (c).

## Discussion

Here, we developed and characterized 3D models of skin and epidermis and demonstrated epidermal pigment dispersion that is comparable to that observed in human skin. Furthermore, detailed TEM analysis revealed ample evidence for the exo/phagocytosis model of melanocore transfer. We obtained images showing exocytosis from melanocytes, melanocores between melanocyte dendrites and keratinocytes, phagocytosis of melanocores by keratinocytes and finally, MKSs bounded by a single membrane.

We started by demonstrating that our RHPS model shows both the stratified layers typical of human epidermis and melanin dispersion throughout the keratinocyte layers, closely representing distinguishing features of human skin. An additional model designated RHPE is also presented herein. Whilst the RHPS model includes a dermis-like layer and can be considered a more complete model system, we have also developed a model that consists of the epidermal layer only upon a collagen-coated membrane support. This RHPE model shows the same stratified layers observed in human skin and in our RHPS but omits the underlying dermis layer containing fibroblasts. Whilst this model therefore lacks the additional extracellular matrix, intercellular communication and complexity contributed by fibroblasts, the resources and time required to form it are reduced. Importantly, the RHPE model shows epidermal pigment dispersal comparable to the full thickness skin model, suggesting that this process is not affected significantly by the lack of dermis in basal conditions. We therefore propose that the newly developed RHPE model could be valuable for studies of skin pigmentation and crosstalk between melanocytes and keratinocytes only.

TEM provided a deeper analysis of the models, revealing characteristic features that facilitate melanin transfer in the epidermis. Firstly, between the layers of keratinocytes, small portions of melanocytes can be identified, which likely represent the dendrites extending from the melanocyte cell body that function to efficiently disperse melanin to multiple keratinocytes *in vivo*. In addition, melanocores were often identified close to melanocyte dendrites, which are likely a result of melanin exocytosis by fusion of melanosomes with the melanocyte plasma membrane. We also observed examples of melanocores that had recently been internalised by keratinocytes, which we believe to be the first published examples of melanocore phagocytosis by keratinocytes shown by TEM. Focussing on keratinocytes, in which melanin is stored, we observed melanin surrounded by a single limiting membrane that is likely derived from the keratinocyte plasma membrane after melanocore internalisation. Our observations by TEM strongly support the exocytosis/phagocytosis model of melanin transfer in the epidermis and are consistent with our previous observations in human skin biopsies (Tarafder et al., 2014).

Melanin transfer remains controversial, and several other mechanisms have been proposed (see Introduction). Notably, evidence suggests that melanin can be transferred by uptake of melanosome-laden vesicles/globules that are shed by melanocytes (Ando et al., 2012). However, in the several specimens of RHPS/RHPE imaged by TEM in this study (3 RHPE and 1 RHPS), we did not identify any examples of the postulated features of other mechanisms, in contrast with the abundant evidence observed towards the melanocore transfer model. We therefore propose that melanocore transfer is the predominant pigment transfer mechanism in the human 3D models presented here.

We note that the epidermal pigmentation observed here is formed in the absence of significant UVr exposure (the models were only exposed to ambient light when removed from the incubator), therefore more closely representing constitutive pigmentation rather than UVr-induced adaptive pigmentation (“tanning”). It is possible that different melanin transfer mechanisms could be more prevalent in different conditions, which we did not explore in the context of this study. Whilst our models are more comprehensive than monoculture or co-culture systems, we note that they do lack further complexities to skin biology *in vivo*, such as the effects of immune cells, variable blood perfusion and mechanical stress. We also acknowledge that artificial skin patches formed *in vitro* using patient-derived induced pluripotent stem cells were already developed, usually for regenerative medicine purposes (Uitto, 2011). However, our model has a notable advantage over these for *in vitro* studies, as it is formed with commercially available primary cells that are fully differentiated, omitting the requirement for technically challenging, costly and failure-prone differentiation protocols.

We propose that these models will be valuable tools for the academic, industrial, pharmaceutical, and cosmetic industries for studies of skin pigmentation control mechanisms. For example, the presented models can be assembled with gene-silenced melanocytes or keratinocytes to study rare pigmentary disorders that show very subtle phenotypes in monoculture or co-culture conditions, but may present clearer phenotypes *in vivo* and in the RHPS/RHPE models over long-term studies. In addition, pharmacological agents can be tested by either addition to the basal media or by application to the apical surface (at the ALI), mimicking systemic and topical application, respectively. Moreover, there is increased demand to reduce the use of animals for experimentation in the research and pharmaceutical industries, and our model could provide an alternative to reduce studies on animals. These factors, combined with the relative ease of formation and availability of resources, highlight the high potential value of this novel model system in skin pigmentation research.

## Acknowledgements

We thank the Electron Microscopy Facility at the Instituto Gulbenkian de Ciência, the Pathological Anatomy service in the Instituto Português de Oncologia de Lisboa Francisco Gentil (IPOLFG), and the Histology Facility at CEDOC for technical assistance. This project was supported by Fundação para a Ciência e a Tecnologia (FCT), Portugal through FCT Unit iNOVA4Health – UID/Multi/04462/2013 co-funded by FEDER under the PT2020 Partnership Agreement, grant PTDC/BIA-CEL/29765/2017 (MJH, DCB), PhD fellowships 2020.08528.BD (SLV), PD/BD/114118/2015, PD/BD/137442/2018 (MVN) and PD/BD/136905/2018 (JC), and the FCT Investigator Program to DCB (IF/00501/2014/CP1252/CT0001).

