## Supplementary Figures for "Reconstructed human pigmented skin/epidermis models achieve epidermal pigmentation through melanocore transfer"

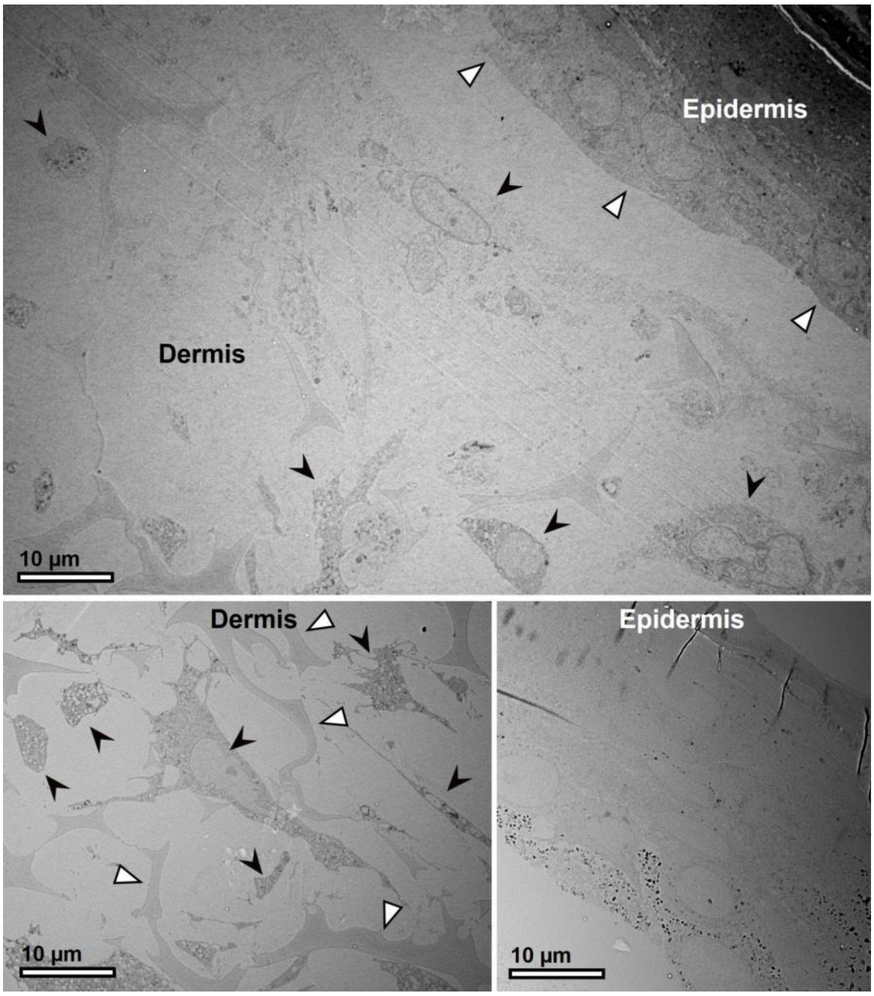


**Figure S1: Reconstructed human pigmented skin visualised by transmission electron microscopy.** A specimen of RHPS observed by TEM shows epidermis upon a support incorporating fibroblasts. The reconstructed epidermis (top panel, white arrowheads) is formed on top of a dermis, formed by fibroblasts (black arrowheads, top and bottom left panel) within a polystyrene scaffold support (white arrowheads, bottom left panel). Scale bar, 10 µm.

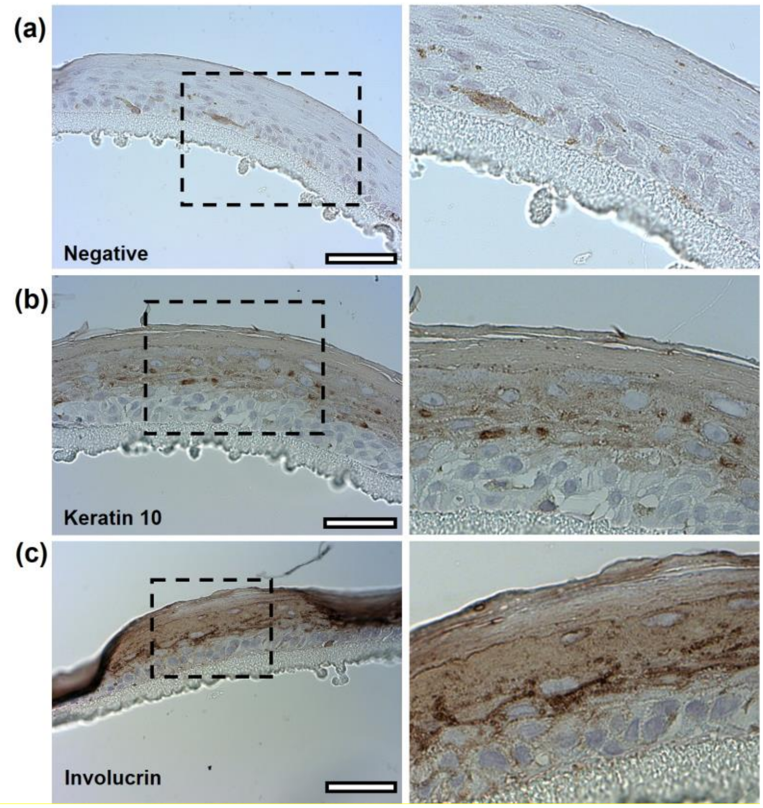


**Figure S2: Immunohistochemistry of epidermal markers in reconstructed human pigmented epidermis model.** Epidermal markers that show the stratification of epidermis in RHPE. A negative control for the staining is shown in **(a)**. Keratin 10 **(b)** stains the suprabasal layers and involucrin **(c)** stains the *stratum spinosum* and *stratum granulosum*. Scale bar, 50 µm.
